# Friend or Foe: Hybrid proline-rich proteins determine how plants interact with and respond to beneficial and pathogenic microbes

**DOI:** 10.1101/2021.08.26.457806

**Authors:** Zeeshan Z. Banday, Nicolás M. Cecchini, Allison T. Scott, Ciara T. Hu, Rachael C. Filzen, Elinam Agbo, Jean T. Greenberg

## Abstract

Plant plastids generate signals, including some derived from lipids, that need to be mobilized to effect signaling. We used informatics to discover potential plastid membrane proteins involved in microbial responses. Among these are proteins co-regulated with the systemic immunity component AZI1, a hybrid proline-rich protein (HyPRP) and HyPRP superfamily members. HyPRPs have a transmembrane domain, a proline-rich region (PRR) and a lipid transfer protein domain. The precise subcellular location(s) and function(s) is unknown for most HyPRP family members. As predicted by informatics, a subset of HyPRPs have a pool of protein that targets plastid outer envelope membranes (OEMs) via a mechanism that requires the PRR. Additionally, two HyPRPs may be associated with thylakoid membranes. Most of the plastid and non-plastid localized family members also have pools that localize to endoplasmic reticulum, plasma membrane or plasmodesmata. HyPRPs with plastid pools regulate, positively or negatively, systemic immunity against the pathogen *Pseudomonas syringae*. HyPRPs also regulate the interaction with the plant growth promoting rhizobacteria *Pseudomonas simiae* WCS417 in the roots to influence colonization, root system architecture and/or biomass. Thus, HyPRPs have broad and distinct roles in immune, development and growth responses to microbes and reside at sites that may facilitate signal molecule transport.

## Introduction

Plants have evolved strategies to provide immunity against a broad range of pathogens. The recognition of pathogen-derived molecules at the surface or inside the cell leads to the establishment of local or systemic immunity (Albert et al., 2020; Fu and Dong, 2013; Zipfel, 2008). During systemic immunity, the host plant responds to a primary infection to provide a faster response against a broad range of secondary infections by a phenomenon called priming (Jaskiewicz et al., 2011; Jung et al., 2009; Pieterse et al., 2014). The recognition of pathogens/pathogen-derived molecules in the leaves induces systemic acquired resistance (SAR), a broad spectrum program that suppresses diverse pathogens at sites distal to a primary infection (Fu and Dong, 2013). Colonization of roots with beneficial bacteria can lead to induced-systemic resistance (ISR) in the aboveground part of the plant (Pieterse et al., 2014). These systemic immune programs require long distance transport of mobile signal(s) from the immunized tissue (Carella, 2020; Cecchini et al., 2019b; Kachroo and Kachroo, 2020; Pieterse et al., 2014). Many candidate signal molecules, including azelaic acid (AZA), have been identified as mobile signals for SAR. These signals may act together to confer SAR (Jung et al., 2009; Kachroo and Kachroo, 2020). Although there are differences in the mechanism of induction, some of the signaling components for SAR and ISR overlap (Cecchini et al., 2019b, 2015; Pieterse et al., 1996, 1998; Beckers and Conrath, 2007; Beckers et al., 2009). Beneficial rhizobacteria not only induce ISR, but also stimulate the growth and productivity of plants and impact the root system architecture (Berendsen et al., 2012; Efthimiadou et al., 2020). A crosstalk between the signaling pathways of rhizosphere-induced growth promotion and ISR remains unknown. However, some degree of shared signaling seems plausible (Bulgarelli et al., 2013; Haney et al., 2015).

Plastids coordinate many cellular responses to infections, generating signals that activate defenses (Bobik and Burch-Smith, 2015). Several key defense molecules including hormones and second messengers are synthesized in plastids (Bobik and Burch-Smith, 2015; Dempsey et al., 2011; Mittler et al., 2004). More importantly, signals needed for SAR, such as salicylic acid, AZA and pipecolic acid are produced at least in part within plastids (Dempsey et al., 2011; Hartmann et al., 2017; Zoeller et al., 2012; Rekhter et al., 2019). These molecules (or others regulated by them) should be mobilized from their site(s) of synthesis for long-distance transport. The SA precursor isochorismate synthesized in the chloroplasts is thought to be exported by ENHANCED DISEASE SUSCEPTIBILITY5 (EDS5) (Rekhter et al., 2019; Serrano et al., 2013). Similarly, pools of AZELAIC ACID INDUCED 1 (AZI1) and its paralog EARLY ARABIDOPSIS ALUMINUM INDUCED 1 (EARLI1) are localized in the plastid envelopes and help in the mobilization of plastid-produced AZA (Cecchini et al., 2015; Zoeller et al., 2012). These proteins also localize to the endoplasmic reticulum (ER), and plasma membrane (PM), including plasmodesmata (PD) (Cecchini et al., 2015). Pathogen infection or treatment with the microbe (pathogen)-associated molecular pattern MAMP/PAMP flg22, an epitope of bacterial flagellin, induces AZI1 and EARLI to become enriched at chloroplasts (Cecchini et al., 2015, 2020). The plastids together with other membranous organelles like nucleus, ER and PM (including PD), facilitate the efficient relay of inter-organellar signaling to achieve whole-cell immunity (Helle et al., 2013; Toulmay and Prinz, 2011; Caplan et al., 2015; Lee, 2015). AZI1 and EARLI1 are essential for SAR, MAMP/PAMP-triggered SAR, ISR and AZA-induced systemic disease resistance (Cecchini et al., 2019b, 2015). AZI1 is a signal-anchored protein that has an N-terminal bipartite signal composed of a signal peptide (SP)-like hydrophobic domain followed by a charged protein region (CPR, consisting of at least three Lys and/or Arg residues, within eight amino acids at the C-terminal side of TMD) and proline rich region (PRR) for the plastid targeting (Cecchini et al., 2015, 2020; Lee et al., 2011). The SP-like region acts as a transmembrane domain (TMD) anchoring AZI1 to the membranes, whereas the PRR guides its plastid association (Cecchini et al., 2020). The superfamily of proteins to which AZI1 belongs are called hybrid proline-rich proteins (HyPRPs). Invariant features include the SP-like region and a C-terminal 8-cysteine motif (8CM)/lipid transfer protein-like (LTP) domain. Most members of the family also have a PRR between these regions that varies in length. Although AZI1 uses a signal-anchored mechanism for plastid targeting, it does not conform to targeting features of classical signal anchor proteins whose N-termini have low hydrophobicity scores on the Wimley-White hydrophobicity scale (Lee et al., 2011).

In this work, we used bioinformatic analyses to identify proteins that, like AZI1, have a pool of protein that localizes to plastid membranes, but which are not classical signal anchored proteins. Included in the screen are proteins whose transcripts are co-regulated with AZI1 and members of the HyPRP superfamily, respectively. Herein, we show the utility of our bioinformatics approach for finding proteins with a pool that localizes to plastid membranes and show that some of them, in the HyPRP superfamily, have different positive or negative roles in systemic immunity, development and growth responses to microbes in roots and shoots.

## Results

### *In silico* analysis of subcellular targeting of the Arabidopsis proteome

Well-known targeting algorithms (TargetP, ChloroP and SignalP; (Emanuelsson et al., 1999, 2000; Nielsen et al., 1997) predict AZI1 to be a secreted protein. However, a pool of AZI1 (and that of close paralogs) localizes to plastid envelopes using an N-terminal bipartite signal (SP-like/TMD + PRR; (Cecchini et al., 2015)) AZI1 has similar properties to apicomplexan proteins that target apicoplasts (non-photosynthetic plastids) using N-terminal bipartite signal sequences (**Figure 1**; (Cecchini et al., 2020)). The PATS algorithm, trained using apicomplexa plastid proteins, can successfully predict plastid localization of AZI1 and paralogs by searching for this bipartite signal (Cecchini et al., 2015, 2020; Zuegge et al., 2001). We used PATS to predict N-terminal bipartite signal sequence and plastid localization of the total Arabidopsis proteome. Overall, the PATS-generated prediction output (positive and negative selections) showed that approximately 7-8% proteins coded by each chromosome may localize to plastids (**Figure 1B**). The Arabidopsis proteins predicted to target to the plastids by PATS are listed in **Supplementary Dataset S1**. Interestingly, the gene ontology (GO) functional characterization (biological processes) of these proteins revealed that among others, the GO terms related to cell communication, response to biotic stimulus, defense response, protein phosphorylation, cell wall organization and cell-cell signaling were significantly overrepresented (**Supplementary Figure S1A**).

**Figure 1.**
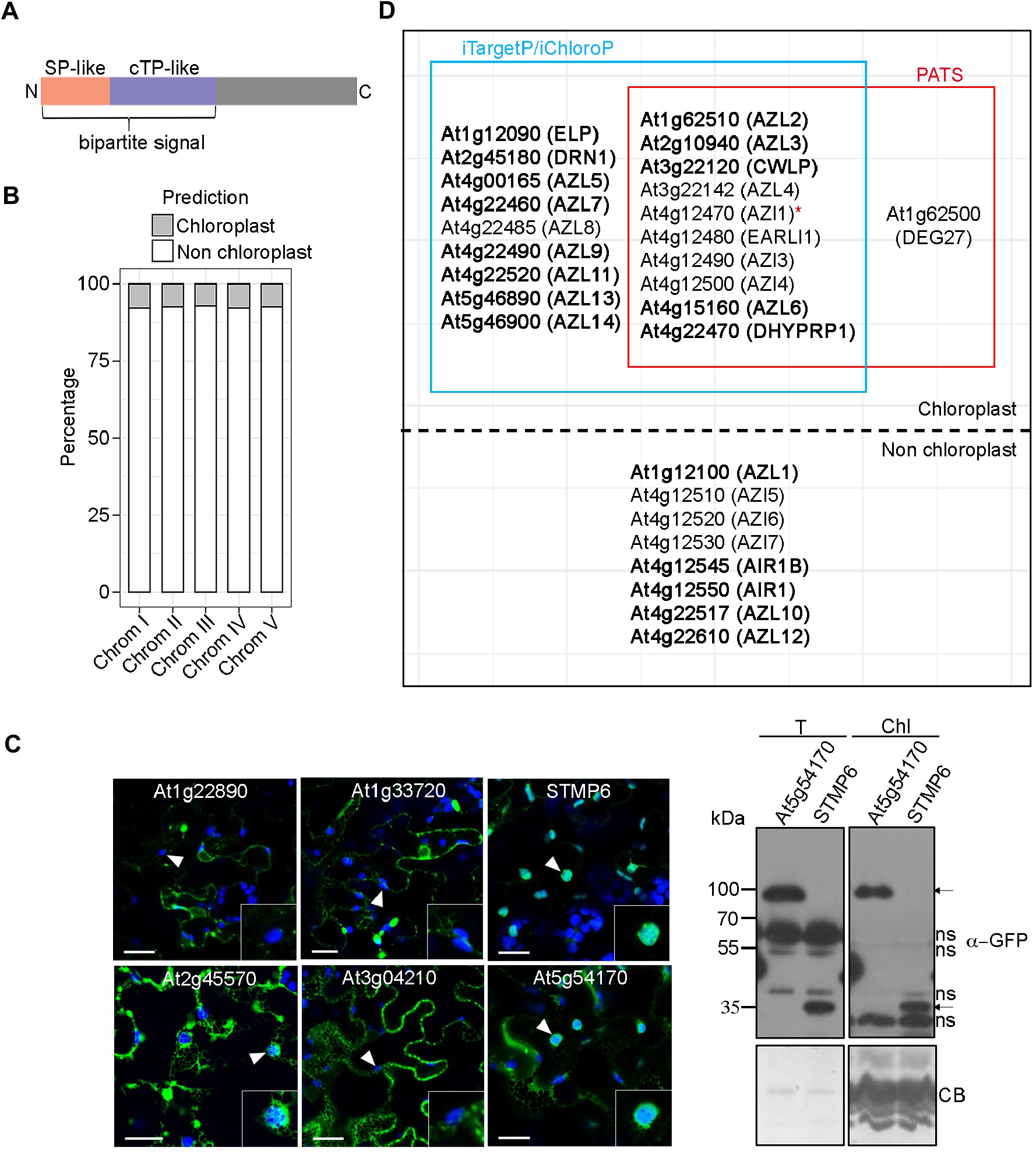
Targeting of Arabidopsi*s* bipartite signal proteins to plastids. (**A**) Schematic of N-terminal bipartite signal of Arabidopsis HyPRPs/Apicomplexan proteins predicted by PATS. SP-like is signal-peptide like and cTP-like is chloroplast transit-peptide like sequence. (**B**) The percentage prediction output of PATS algorithm for each chromosome of Arabidopsis. The full-length sequences of Arabidopsis proteome (ATpepTAIR10) were analyzed by PATS for the predicted localization (%) to plastids. (**C**) Left panels: Laser scanning confocal micrographs of epidermal cells showing localization of non-HyPRP:GFP proteins encoded by the indicated genes transiently expressed in *N. benthamiana* leaves under Dex control. Samples were imaged 21-24h after infiltration with 30µM Dex. White arrowheads indicate chloroplasts (shown in insets). Scale bar is 20µm. Right Panel: Immunoblots of total (T) and chloroplast (Chl) fractions of *N. benthamiana* leaves transiently expressing At5g54170-encoded and STMP6 GFP-tagged proteins. Chloroplast fractions were isolated from leaves 21h after treatment with 30µM Dex. Signal was observed using anti-GFP antibody. ns is non-specific band; arrows indicate specific band. Coomassie blue (CB) staining shows the protein loading. (**D**) Targeting prediction of Arabidopsis HyPRPs by iterative TargetP/ChloroP and PATS algorithms. For iTargetP/iChloroP, the sequences without N-terminal SP-like region were used. The proteins with targeting studied herein are in bold. The asterisk indicates AZI1, a variant of signal-anchored protein, where SP-like domain is a transmembrane (TMD) (Cecchini et al., 2020). HyPRPs not predicted by either method to be plastid-localized are in the lower section of the panel.

To test our predictions, we selected six PATS-positive proteins whose transcripts showed similar expression profiles and clustered together with AZI1 as assessed by the GENVESTIGATOR^®^ Hierarchical clustering tool (biological context: Perturbations; (Hruz et al., 2008)). As a secondary screen, we chose proteins that did not conform to a key characteristic of classical signal-anchored plastid OEM proteins. Specifically, their SP-like N-terminal regions lacked low hydrophobicity scores (<0.4) on the Wimley-White hydrophobicity scale (Lee et al., 2011). Instead, the scores (calculated with the MPEx algorithm (http://blanco.biomol.uci.edu/mpex; (Jayasinghe et al., 2001; Snider et al., 2009)) were characteristic of ER-targeted proteins (>0.4): AT5g54170: 0.58, AT1G22890/STMP2: 0.63, AT2G45570.1/CYP76C2: 0.90, AT1G65500/STMP6: 0.63, AT1G33720/CYP76C6: 0.81, AT3G04210.1/TN13: 0.50. We then assessed protein localization by confocal microscopy of C-terminal GFP fusions expressed under dexamethasone (Dex) control in *Nicotiana benthamiana*. Three candidates encoded by At5g54170, At1g65500/*STMP6* and At2g45570, respectively, showed GFP signals that co-localized to varying extents with plastid signals (**Figure 1C, Left panel)**. The subcellular locations of the At5g54170-encoded protein and STMP6 were further corroborated by fractionation and immunoblotting **(Figure 1C, Right panel)**. This shows that the PATS algorithm can identify some plastid proteins besides AZI1 and EARLI1 from Arabidopsis.

### *In silico* analysis of subcellular targeting of the Arabidopsis HyPRPs

PATS also predicted that seven HyPRP members, besides AZI1 and close paralogs, are plastid localized (**Figure 1D**). Arabidopsis HyPRPs are present in gene clusters in the genome (**Supplementary Figure S1B**). Some members of the HyPRPs family were previously named (AZI1, EARLI1, AZI3-AZI7, DEG27, DRN1, DHyPRP1, AIR1, AIR1B, ELP and CWLP); we named the remaining HyPRPs as AZL (<u>AZ</u>I1-<u>L</u>IKE) (**Figure 1D**). Considering that the PATS algorithm was trained using apicomplexa proteins, we also implemented another approach to identify AZI1-like plastid-targeting signals in Arabidopsis HyPRPs. TargetP and SignalP can efficiently recognize SP-like sequences and their cleavage sites in plant proteins. Therefore, we manually removed the N-terminal SP-like sequences identified by these algorithms. These SP-like-minus sequences of HyPRPs were then reanalyzed by TargetP and ChloroP (referred to herein as iterative TargetP, iTargetP or iChloroP) customized for plants with default cutoffs. By using SP-like-depleted sequences, the algorithms can analyze the N-terminal transit peptide-like sequences (called cTPs) for targeting specificity. About 70% of the Arabidopsis HyPRPs were predicted to localize to plastids (**Figure 1D, Supplementary Table S1**). Moreover, the PATS output had a high overlap with that of iTargetP/iChloroP, suggesting that apicoplast or iterative chloroplast predictors can be successfully deployed to find candidates of plant plastid membrane proteins with SP-like motifs.

### Arabidopsis HyPRPs localize to the expected sites for the defense signal mobilization

The expected sites of defense signal movement are plastids, ER, PM including PD, and the multiple membrane contact sites formed by them (Cecchini et al., 2015; Lee, 2015; Lim et al., 2016). Using confocal microscopy, eighteen HyPRP-GFP fusion proteins produced in *N. benthamiana* were found to target the expected sites of defense signal transport (**Figure 2**). Based on comparison with our previous study of AZI1 localization (Cecchini et al., 2015), these patterns of GFP signals could be grouped as: (1) ER and PM including foci similar to PD (AIR1, AIR1B, AZL1, AZL5, AZL7, AZL9, AZL10, AZL11, AZL12), (2) chloroplasts plus ER (AZL13), and (3) chloroplasts plus ER and PM including foci similar to PD (ELP, AZL2, CWLP and AZL14). Initially, we cautiously assigned DRN1 to group three, because although there were some chloroplast signals in the micrographs (**Figure 2A**), in many cases GFP was detected in nuclei (**Supplementary Figure S1C**). This raised the possibility that DRN1-GFP was proteolyzed. Indeed, GFP was cleaved from a large pool of the protein **(Supplementary Figure S1C)**. Two previous mass spectrometry experiments assigned DRN1 to plastid membranes (Peltier et al., 2004; Tomizioli et al., 2014), providing additional confidence that the protein is plastid-associated.

**Figure 2.**
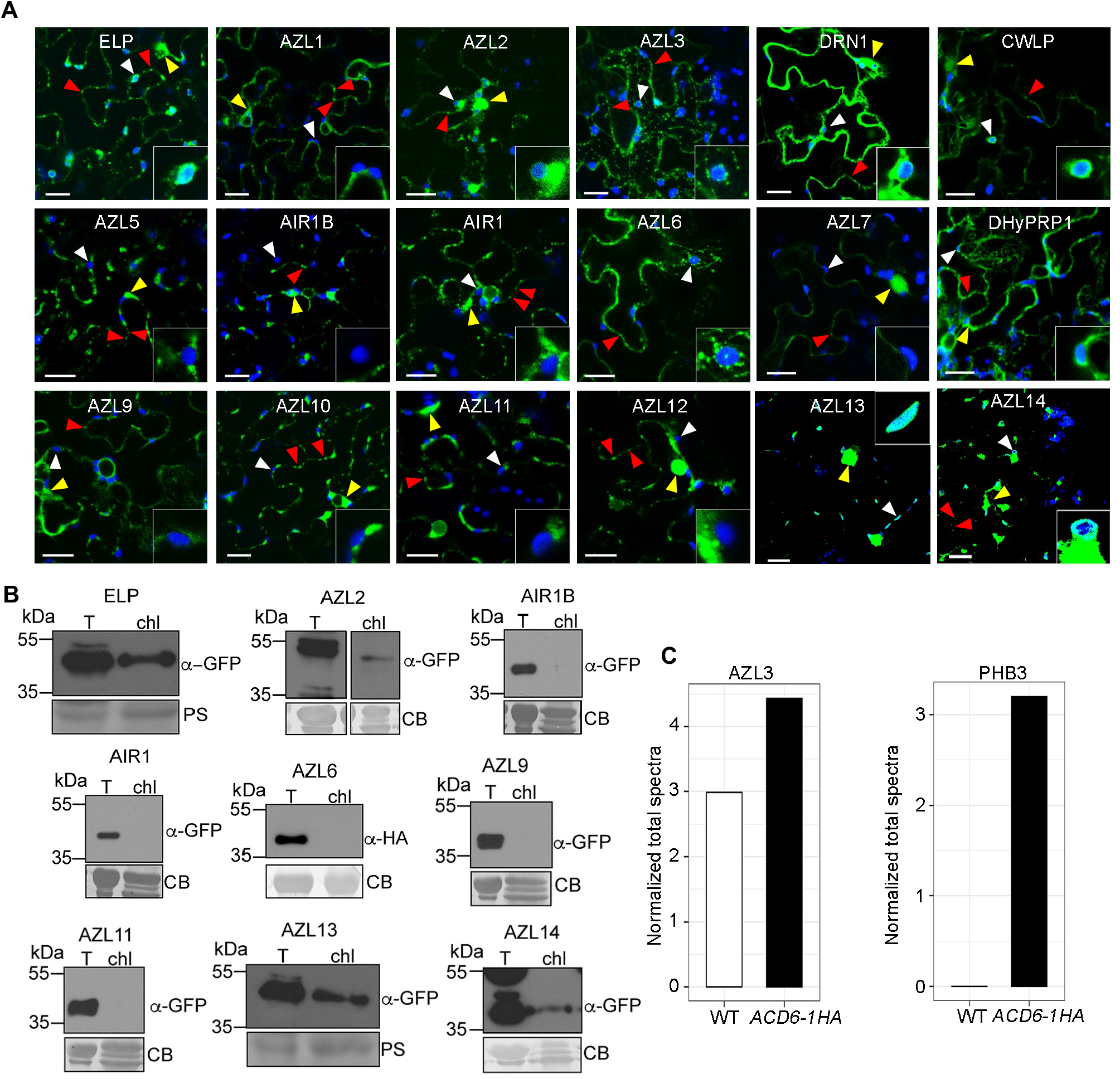
Subcellular localization of HyPRP:GFPs in *N. benthamiana*. (**A**) Laser scanning confocal micrographs of epidermal cells showing localization of various HyPRP:GFP proteins transiently expressed in *N. benthamiana* leaves under Dex-inducible promoter. Samples were imaged 21-24h after infiltration with 30µM Dex. White arrowheads indicate chloroplasts (shown in insets), red arrowheads show plasmodesmata-like structures/plasma membrane and yellow arrowheads show dense GFP signal in perinuclear ER. Scale bars are 20µm. GFP, green; chloroplast autofluorescence, blue. (**B**) Immunoblots of total (T) and chloroplast (chl) fractions of *N. benthamiana* leaves transiently expressing a subset of HyPRP:GFP proteins shown in (**A**). Chloroplast fractions were isolated from leaves 21h after treatment with 30µM Dex. Signal was observed using anti-GFP or anti-HA antibody. Coomassie blue (CB) or Ponceau S (PS) staining shows the protein loading. (**C**) Normalized total spectra of peptides matching AZL3 in chloroplast membranes of constitutively defense active *ACD6-1HA* and WT. PHB3, a known chloroplast envelope protein is shown as control (Seguel et al., 2018). Total spectra normalization was analyzed by Scaffold software for each sample (WT or *ACD6-1HA*).

Generally, the chloroplast-targeted proteins showed a ring-like pattern of fluorescent signal around them. In one case (AZL3), punctate GFP signals around many chloroplasts in addition to PM signals were observed. While AZL6 showed similar localization to AZL3, the chloroplast-associated punctate signals were less prominent. Finally, DHYPRP1 showed ER, PM/PD and incomplete chloroplast rings (**Figure 2A**). The patterns showing ER, PM/PD and chloroplast localization of HyPRPs were largely similar to the family member AZI1 and close paralogs (Cecchini et al., 2015, 2020).

Evaluation by immunoblotting of chloroplast fractions from *N. benthamiana* leaves transiently expressing many HyPRP-GFP fusion proteins largely corroborated the microscopy (**Figure 2B**). We were unsuccessful in using immunoblotting to detect AZL3-GFP after fractionation due to interference from the rubisco large subunit. Instead, we used a proteomics approach and found AZL3 to be differentially enriched in chloroplast membranes of a constitutively defense active Arabidopsis line (*ACD6-1HA*, (Lu et al., 2005)) relative to WT (**Figure 2C**). Prior proteomics studies detected AZL3 in the thylakoid and envelope fractions, respectively (Tomizioli et al., 2014; Kleffmann et al., 2004).

We next analyzed the localization patterns of Dex-inducible GFP fusions of the HyPRPs AZL3, AZL13, AZL14 and ELP in stable Arabidopsis transgenic lines. Each transgene was expressed in the respective mutant background. A portion of the GFP signal of these four fusion proteins colocalized with chloroplast autofluorescence (**Figure 3**). Interestingly, while the GFP signals for AZL13, AZL14 and ELP each showed a ring-like pattern surrounding the chloroplasts, AZL3 had a more diffuse signal. This was further confirmed by analyzing the fluorescence intensity profiles of the merged images (**Figure 3, right most panel**) and is consistent with its import and association with thylakoid membranes (Tomizioli et al., 2014). Inspection of the features of the PRR for AZL3 compared with other HyPRPs that show the ring-like pattern revealed that the AZL3 PRR lacks a KP motif present in others and also possesses a unique repeated sequence (**Supplementary Table S1**). These and/or other differences in the PRR composition (e.g. abundance of prolines relative to charged amino acid residues) may account for differences in the suborganellar plastid localization patterns. Altogether, our data show that for several HyPRPs, a pool of these proteins localizes to chloroplasts, as predicted.

**Figure 3.**
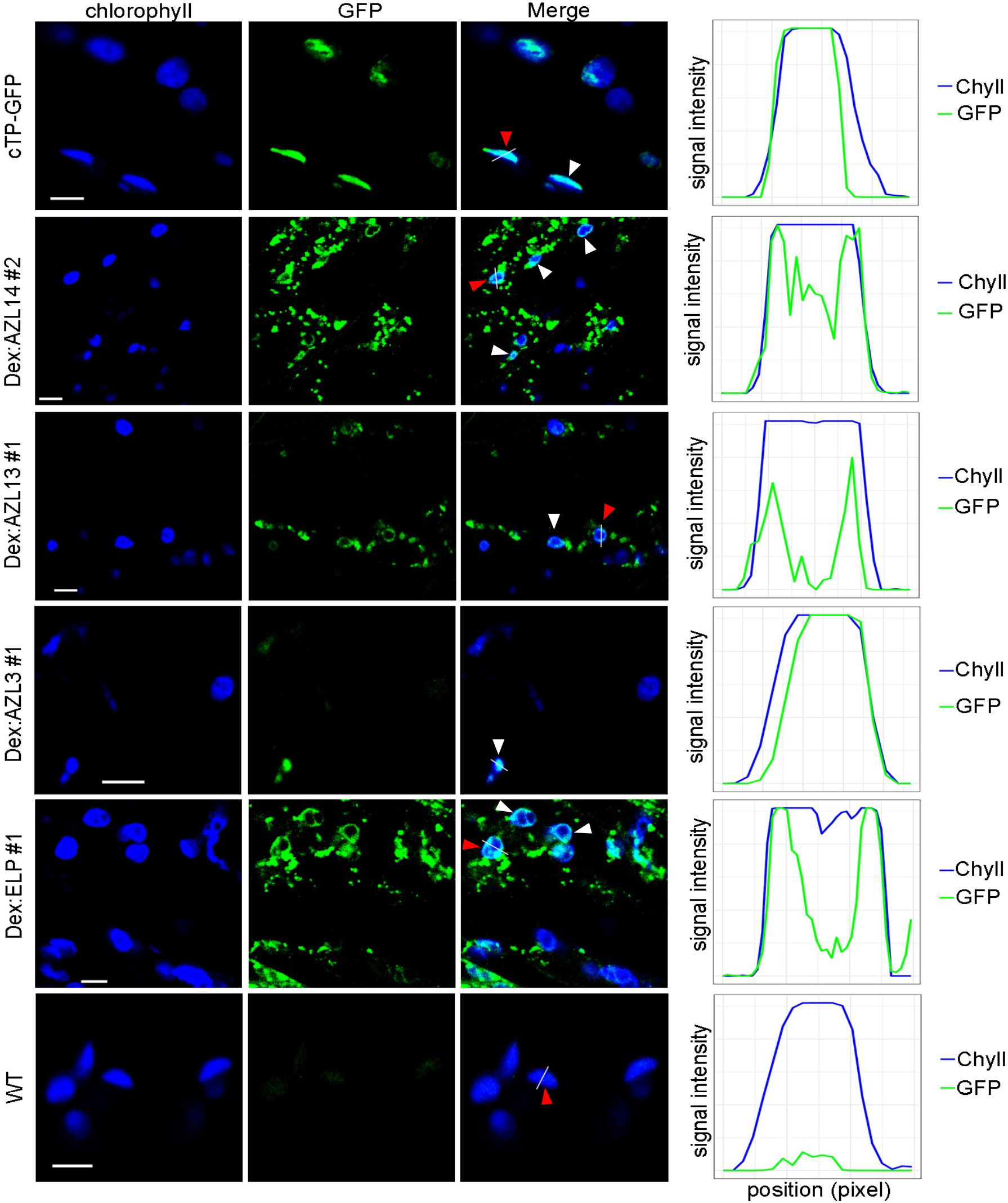
Chloroplast localization of HyPRP:GFP proteins in Arabidopsis transgenic lines. Laser scanning confocal micrographs of epidermal cells showing chloroplast localization of indicated GFP tagged HyPRPs in 9-12 day-old Arabidopsis transgenic seedlings (hypocotyl). Plants were sprayed with 30µM Dex plus 0.04% Tween 20, 21-24h prior to imaging. Fluorescence intensity profiles in the merged images show the overlap of GFP (green) and chlorophyll (Chyll, blue) signal intensities along solid white lines (indicated by the red arrowheads). Note the overlap of GFP (green) and chlorophyll (Chyll, blue) intensities along chloroplast peripheries. The plastid-targeted GFP line (cTP-GFP) (Jiang et al., 2021) and untransformed WT (Col-0) serve as positive and negative controls, respectively, for this experiment. The images were analyzed by ImageJ (Fiji). Similar results were observed in two independent experiments. White arrowheads indicate chloroplast localization. Scale bar is 5µm. GFP, green; chloroplast autofluorescence, blue.

### Several chloroplast-localized HyPRPs are associated with OEMs

Except for AZL1 and AZL9, N-terminal regions of all HyPRPs have hydrophobicity scores >0.4 on the Wimley-White scale (**Supplementary Table S1**). Thus, they do not have features of classical signal-anchored OEM plastid proteins (Lee et al., 2011). Moreover, AZL1 and AZL9 (TMD hydrophobicity < 0.4; plastidial score) did not target to the chloroplasts as indicated by confocal microscopy (**Figure 2A**). This suggests that HyPRPs follow a non-canonical mechanism of targeting to chloroplast membranes, similar to the family member AZI1 (Cecchini et al., 2020).

To study the above possibility, we analyzed the chloroplast membrane association of HyPRPs that showed ring-like localization around chloroplasts. We isolated chloroplasts from *N. benthamiana* leaves transiently expressing GFP-tagged AZL2, AZL13, AZL14 and ELP. The chloroplasts were then partitioned into pellet (membrane) and soluble fractions. Interestingly, all these GFP-tagged HyPRPs were found to be associated with pellets (chloroplast membranes) as evaluated by immunoblotting (**Figure 4**). Furthermore, intact chloroplasts from plants transiently producing HyPRP-GFP fusion proteins treated with thermolysin resulted in HyPRP-GFP proteolysis, consistent with localization to the OEMs. In contrast, the intermembrane space-located POTRA domains of OEM protein Toc75 and stromal HSP70 were fully protected from thermolysin digestion (**Figure 4B**; (Chen et al., 2016; Seguel et al., 2018)). These experiments indicate that several chloroplast-targeted HyPRPs reside in OEMs and that their C-terminal regions topologically face the cytosol.

**Figure 4.**
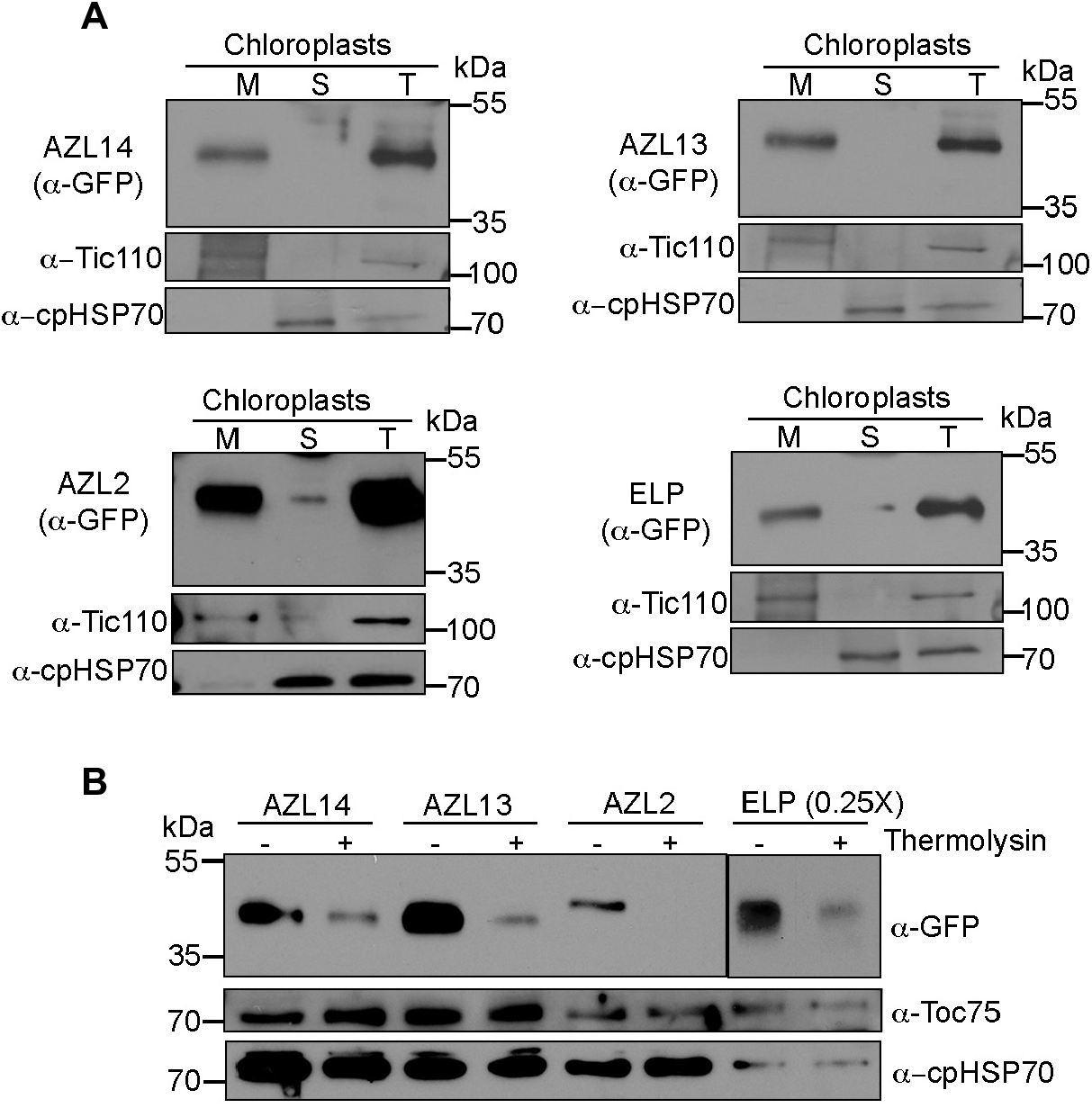
Chloroplast OEM localization of Arabidopsis HyPRPs in N. *benthamiana*. (**A**) Immunoblots of total (T), membrane (M) and soluble (S) fractions of isolated chloroplasts from *N. benthamiana* leaves transiently expressing AZL14, AZL13, AZL2 and ELP tagged with GFP at C-terminal under Dex-inducible promoter. HyPRPs were detected with ⍺-GFP antibody. ⍺-Tic110 and ⍺-cpHSP70 antibodies were used to detect the markers of inner envelope membrane, Tic110 and stroma, HSP70. (**B**) Immunoblots of intact chloroplast fractions expressing HyPRP:GFPs as in (**A**) treated with buffer (-) or thermolysin (+). The digestion of HyPRP:GFPs by thermolysin was evaluated by ⍺-GFP antibody. Due to the higher observed expression of ELP:GFP compared to other HyPRP:GFPs, 1/4X total protein was loaded for this sample. A low exposure for ELP:GFP with ⍺-GFP antibody is shown. The antibodies for OEM protein Toc75 and stromal HSP70 were used to detect the non-digestible controls. These experiments were repeated two-three times with similar results.

### The proline-rich region is required for the chloroplast targeting of HyPRPs

Multiple sequence alignment of Arabidopsis HyPRPs revealed conserved N-terminal SP-like hydrophobic and C-terminal 8CM/LTP domains, and highly variable PRRs, variable both in terms of the number of Pro residues and the length of each PRR (**Supplementary Figure S2, Supplementary Table S1**). AZI1 uses a non-canonical signal-anchored targeting mechanism that depends on the PRR for a pool of AZI1 to target chloroplasts (Cecchini et al., 2020). To determine whether this is true for other members, we deleted the PRRs (**Figure 5**) of AZL2, AZL13, AZL14, and ELP. We observed GFP cleavage for AZL14ΔPRR (AZL14^Δ26-44^), which was circumvented by deleting a portion of its LTP together with the PRR (AZL14^Δ24-101^). The deletion of PRRs prevented all these HyPRP-GFP fusion proteins from targeting to chloroplasts, consistent with a role for the PRRs in promoting the association of some Arabidopsis HyPRPs with chloroplasts (**Figure 5A-C**).

**Figure 5.**
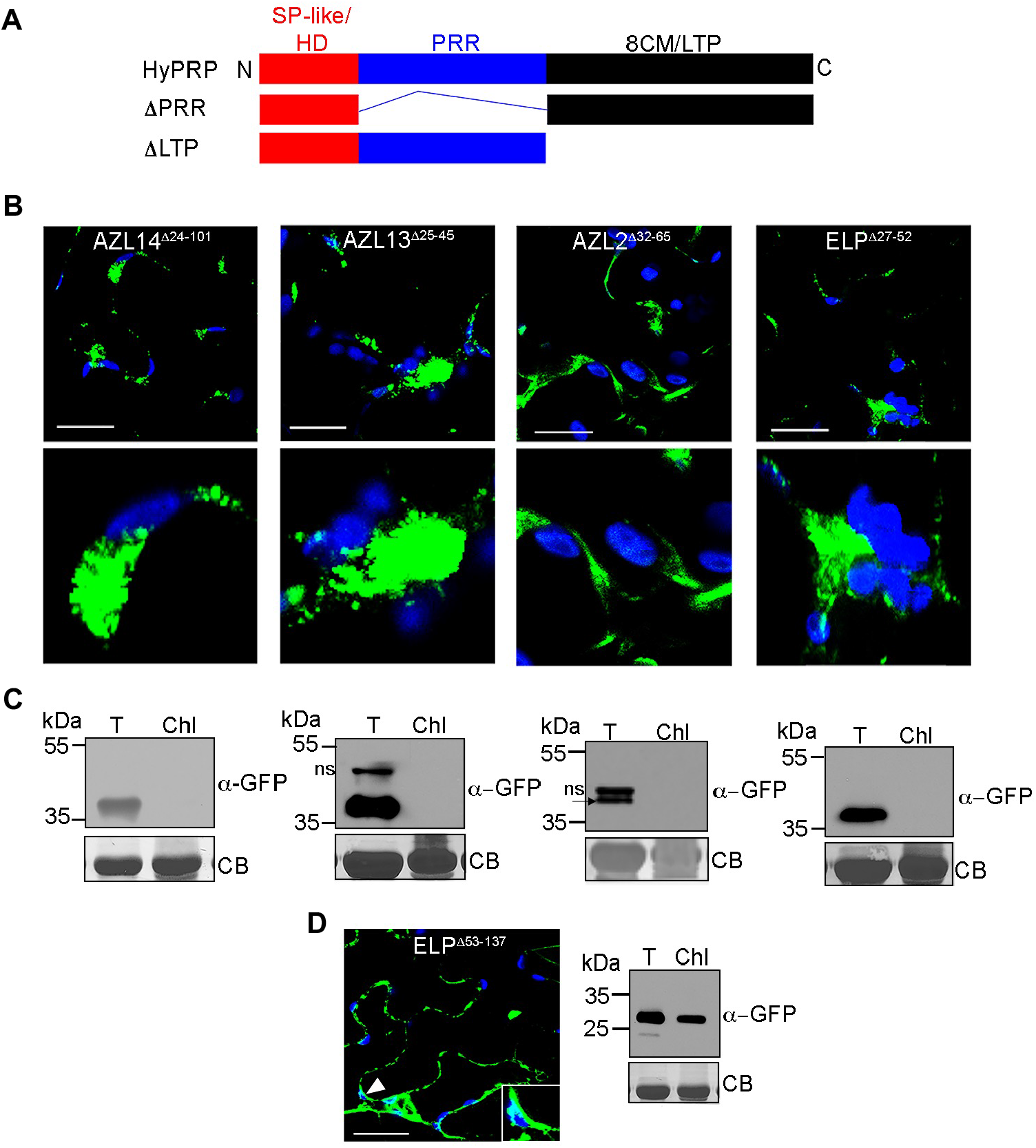
Subcellular localization of the truncated versions of HyPRP:GFPs in *N. benthamiana*. (**A**) Schematic of a HyPRP showing N-terminal SP-like or HD, PRR and the C-terminal 8CM/LTP domain. The ΔPRR and ΔLTP versions are also depicted. (**B**) Confocal micrographs showing the localization of ΔPRR mutants of GFP-tagged HyPRPs in epidermal cells under Dex-inducible promoter. Lower panels show the enlarged views of chloroplasts. (**C**) Immunoblots of total (T) and chloroplast (Chl) fractions from *N. benthamiana* leaves transiently expressing HyPRP:GFP proteins as in (**B**). GFP-tagged HyPRPs were detected by ⍺-GFP antibody. ns is non-specific band; arrow indicates specific band. The same blots stained with Coomassie blue (CB) show protein loading. Similar results were observed in two or more independent experiments for AZL14, AZL13 and ELP. For AZL2, the result of one experiment is shown. (**D**) Confocal micrograph of epidermal cells and immunoblot of total (T) and chloroplast (Chl) fraction showing the subcellular localization of GFP-tagged ELP:ΔLTP. Inset shows signal associated with chloroplast. For the micrographs, GFP is green; chloroplast autofluorescence is blue. All scale bars are 20µm.

In AZI1 and the closely related paralogs EARLI1 and AZI3, the C-terminal 8CM/LTP domains did not contribute to chloroplast targeting (Cecchini et al., 2015, 2020). To check if this domain is similarly dispensable for chloroplast targeting of a more distantly related HyPRP, we generated ELPΔLTP-GFP. This truncated version of ELP associated with chloroplasts (**Figure 5D**). These data show that the PRRs play key roles in chloroplast targeting and are consistent with the N-terminal region of HyPRPs being sufficient to confer this targeting.

### Expression of *HyPRPs* is regulated in response to microbes

The HyPRPs AZI1, EARLI1 and DRN1 are required for defense against pathogens and their transcripts are modulated during infection (Cecchini et al., 2015; Dhar et al., 2020). Mining of the literature and public data indicated that many *HyPRP* transcripts are differentially regulated by pathogens or pathogen-derived molecules (e.g. the PAMP flg22), suggesting that they may play a role during plant defense responses (**Table 1**). Moreover, the expression of many *HyPRPs* was also responsive to inoculation of roots with the beneficial microbe *P. simiae* WCS417 (Stringlis et al., 2018). Analysis of publicly available gene expression data also revealed that a number of HyPRPs are co-regulated to different degrees, suggesting they may be involved in the same processes (**Figure 6**). The HyPRPs with the highest co-regulation values were encoded by genes with strong chromosomal linkage **(Supplementary Figure S1B)**.

**Table 1.** Expression changes of Arabidopsis *HyPRPs* to WCS417, pathogens and PAMPs *CalCuV* is *Cabbage leaf curl virus, Psm is P. syrinage pv maculicola ES4326, Pst* is *P. syringae pv tomato* DC3000, WCS417 is *P. simiae* WCS417, *B. cinerea* is *Botrytis cinerea*, PAMPs is pathogen-associated molecular patterns, *P. brassiccae is Plasmodiophora brassicae*, flg22-PA is flg22 from *P. aeruginosa*, NLP is (Nep1)-like protein

| HyPRP | Pathogen/PAMP/Beneficial microbe-related | References |
| --- | --- | --- |
| At1g12090 (ELP) | *CalCuV* ↓, *Pst* ↓, PAMPs↓, *Psm* ↓, flg22↓(roots) | (Ascencio-Ibáñez et al., 2008; Baum et al., 2019; Stringlis et al., 2018; Thilmony et al., 2006) |
| At1g12100 (AZL1) | flg22↓(roots) | (Stringlis et al., 2018) |
| At1g62500 (DEG27) | WCS417↑(roots), flg22↓(roots) | (Stringlis et al., 2018) |
| At1g62510 (AZL2) | *CalCuV* ↓, *Pst* ↓, PAMPs↓, *P. brassicae* ↑, WCS417↓ (roots), flg22 ↓ (roots) | (Ascencio-Ibáñez et al., 2008; Siemens et al., 2006; Stringlis et al., 2018; Thilmony et al., 2006; Zamioudis et al., 2014) |
| At2g10940 (AZL3) | *CalCuV* ↓, *Pst* ↓, PAMPs↓, *Psm* ↓, WCS417↓ (roots), flg22↓(roots) | (Ascencio-Ibáñez et al., 2008; Baum et al., 2019; Stringlis et al., 2018; Thilmony et al., 2006) |
| At2g45180 (DRN1) | *CalCuV* ↓, *Pst* ↓, PAMPs↓, *Psm* ↓, *B. cinerea* ↓, flg22↓(roots) | (Ascencio-Ibáñez et al., 2008; Dhar et al., 2020; Stringlis et al., 2018; Thilmony et al., 2006) |
| At3g22120 (CWLP) | *CalCuV* ↓, PAMPs↓, *Pst* DC3000/avrRpt2↓ | (Ascencio-Ibáñez et al., 2008; Thilmony et al., 2006; Mine et al., 2018) |
| At3g22142 (AZL4) | WCS417↓ (roots), flg22↓(roots) | (Stringlis et al., 2018) |
| At4g00165 (AZL5) | WCS417↓ (roots), flg22↓(roots) | (Stringlis et al., 2018) |
| At4g12470 (AZI1) | *Psm* ↑, PAMPs ↑, WCS417↑ (roots) flg22PA ↑(roots) | (Cecchini et al., 2020; Jung et al., 2009; Stringlis et al., 2018; Zamioudis et al., 2014) |
| At4g12480 (EARLI1) | *CalCuV* ↑, *Pst* ↑, PAMPs ↑ flg22PA ↑ (1h roots) | (Ascencio-Ibáñez et al., 2008; Gupta and Senthil-Kumar, 2017; Stringlis et al., 2018; Thilmony et al., 2006) |
| At4g12490 (AZI3) | *CalCuV* ↑, PAMPs ↑, WCS417↑(roots), flg22 ↑(roots) | (Ascencio-Ibáñez et al., 2008; Stringlis et al., 2018; Thilmony et al., 2006; Zamioudis et al., 2014) |
| At4g12500 (AZI4) | PAMPs ↑, *Pst* ↑, flg22 ↑, Oomycete NLP↑ | (Mohr and Cahill, 2007; Qutob et al., 2006; Thilmony et al., 2006) |
| At4g12510 (AZI5) | *CalCuV* ↓, WCS417↓ (roots), flg22↓ (roots) | (Ascencio-Ibáñez et al., 2008; Stringlis et al., 2018) |
| At4g12520 (AZI6) | WCS417↓ (roots), flg22↓ (roots), Chitin ↓ (roots) | (Stringlis et al., 2018) |
| At4g12530 (AZI7) | *Pst*↓ | (Howard et al., 2013) |
| At4g12545 (AIR1B) | flg22↓ (roots), Chitin↓(roots) | (Stringlis et al., 2018) |
| At4g12550 (AIR1) | flg22↓(roots), Chitin ↓ (roots) | (Stringlis et al., 2018) |
| At4g15160 (AZL6) | flg22↓ (roots) | (Stringlis et al., 2018) |
| At4g22460 (AZL7) | WCS417↓ (roots), flg22↓ (roots) | (Stringlis et al., 2018) |
| At4g22470 (DHYPRP1) | *Pst* ↑, flg22 ↑, *WCS417* ↑ (roots), flg22 ↑(roots), flg22-PA ↑(roots), Chitin ↑(roots) | (Li et al., 2014; Mészáros et al., 2006; Stringlis et al., 2018) |
| At4g22485 (AZL8) | No known change |  |
| At4g22490 (AZL9) | *Pst* ↑ | (Gupta and Senthil-Kumar, 2017) |
| At4g22517 (AZL10) | No known change |  |
| At4g22520 (AZL11) | No known change |  |
| At4g22610 (AZL12) | *WCS417* ↓ (roots), flg22 ↓ (roots), Chitin ↑(roots) | (Stringlis et al., 2018) |
| At5g46890 (AZL13) | *WCS417* ↓ (roots), flg22-417 ↓ (roots), flg22-PA ↓ (roots) | (Stringlis et al., 2018) |
| At5g46900 (AZL14) | *WCS417* ↓ (roots), flg22-417 ↓(roots), flg22-PA ↓ (roots) | (Stringlis et al., 2018) |

**Figure 6.**
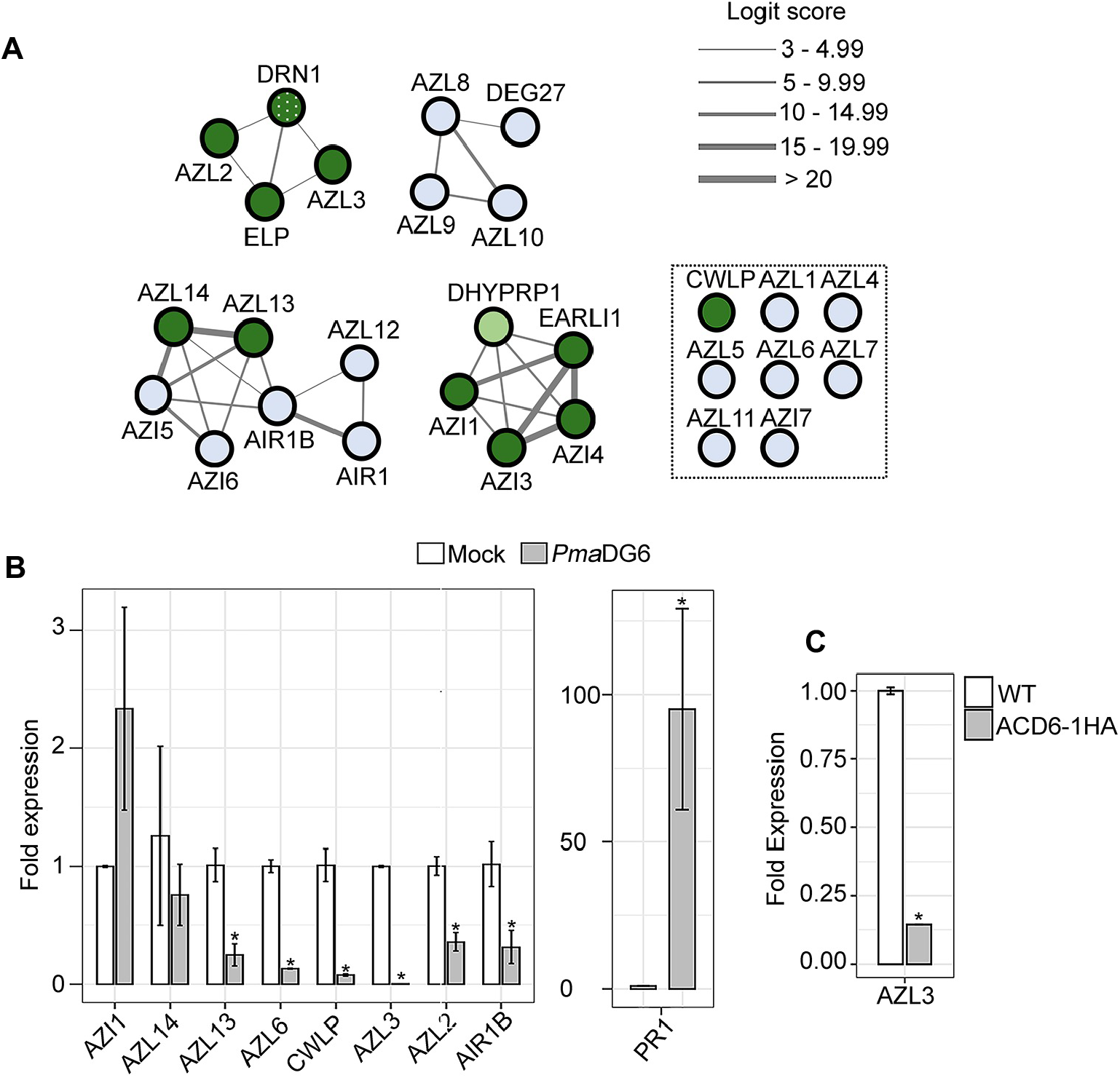
Expression, predicted co-expression and plastid envelope localization of *HyPRPs.* **(A)** Network co-expression map of the *HyPRP* genes using ATTED-II (version 11.0; https://atted.jp/). Solid lines indicate co-expression between the genes included within the top 20 with higher score (Logit Score). Line thickness indicates the co-expression score. Condition-independent and condition-specific co-expression data is depicted. Green circles indicate HyPRPs targeted to the plastid envelope found in this and our previous (Cecchini et al., 2015) study. Dotted green circle indicates DRN1 plastid targeting evidenced in this and other studies (Peltier et al., 2004; Tomizioli et al., 2014). Light green circles indicate partial targeting to plastids. Inside the dashed box are the genes with no co-expression or a Logit score lower than 3. (B) Transcript levels of *HyPRPs* in *Pma*DG6-infected WT (Col-0) leaves 18hpi. PR1 induction shows that the defense response was activated in these plants. (**C**) Transcript levels of AZL3 in constitutively defense active mutant *ACD6-1HA*. In (**B** and **C**) *EF1⍺* was used as an internal reference. Error bars indicate SEM from two biological replicates and two technical replicates. *p<0.05, student’s t-test.

We analyzed the transcript levels of several *HyPRPs* in WT after infection with a pathogen capable of inducing strong local and systemic defenses, *Pseudomonas cannabina pv. alisalensis* carrying AvrRpt2 (avirulent strain *Pma*DG6). Most *HyPRP* transcripts assayed were downregulated by 18h post infection (**Figure 6B**). Because AZL3 showed increased protein levels in the plastid membrane fraction of the constitutively defense-active line ACD6-1HA (**Figure 2C)**, we also assayed its transcript levels in these plants. Interestingly, the AZL3 transcript level was reduced in the ACD6-1HA line relative to WT (**Figure 6C**). Thus, AZL3 may be subject to post transcriptional regulation.

Together, previously published and our data show that the HyPRPs are differentially regulated during biotic interactions (or simulated infection conditions provided by the ACD6-1HA line) and thus may play a role, positive or negative, during interactions with microbes.

### Several HyPRPs regulate SAR in Arabidopsis

Since AZI1 and EARLI1 are required for systemic immunity, other HyPRPs with chloroplast-associated pools might be similarly required. To test this, we evaluated local foliar pathogen resistance and SAR using *azl3, azl13, azl14* and *elp* T-DNA mutants. We used an anti-sense line *AZL2-AS* (Jülke and Ludwig-Müller, 2015) for SAR experiments, since no *azl2* mutant was available.

In all mutants tested, the local (basal) resistance against a SAR-inducing (avirulent) as well as a virulent *Pseudomonas* strain (*Pma*DG6 and *Pma*DG3, SAR-inducing and virulent *Pseudomonas cannabina pv. alisalensis,* respectively) was similar to WT (**Supplementary Figure S3**). In contrast, the *azl3* and *azl13* mutants and the AZL2-AS plants were SAR-deficient, whereas the *azl14* mutant still showed robust SAR, similar to WT (**Figure 7**). SAR response gain values (Jiang et al., 2021) of the different genotypes supported this conclusion (**Figure 7B, D**). Interestingly, the *elp* mutant showed significantly higher resistance levels to *Pma*DG3 in systemic leaves during SAR and an increased response gain compared to the WT (**Figure 7A, B**). This unusual “super-SAR” phenotype was confirmed in *elp-2*, another T-DNA allele (**Supplementary Figure S4A**).

**Figure 7.**
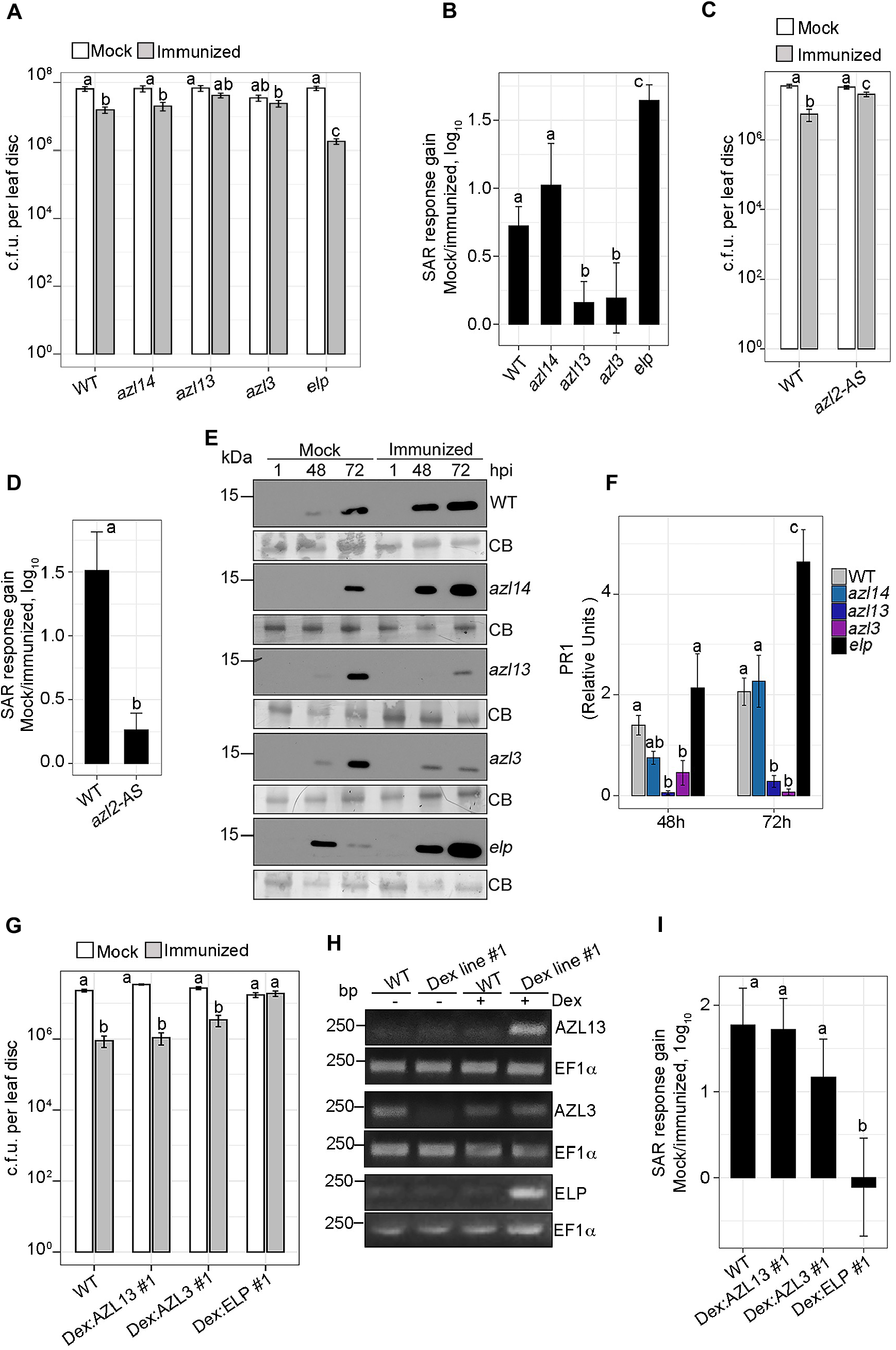
SAR in Arabidopsis *HyPRP* mutants. (**A**, **C**) Growth of virulent bacteria *Pma*DG3 in systemic leaves of WT (Col-0) and indicated mutants three days after infection. The plants were previously immunized in local leaves with avirulent *Pma*DG6 or mock treated, 2 days prior to the secondary infection. The mean c.f.u. of two-three independent experiments are plotted. (**B**) Response gain of SAR due to *Pma*DG6 immunization in (**A**). (**D**) Response gain of SAR due to *Pma*DG6 immunization in (**C**). (**E**) PR1 induction at the indicated time points in the uninfected systemic leaves of plants that were locally mock treated or immunized with *Pma*DG6. (**F**) Densitometric quantification of PR1 levels in immunized samples shown in (**E**). The PR1 levels are relative to the non-specific band in Coomassie blue (CB) stained membrane. The mean from two replicates is shown. Quantification was done by ImageJ (Fiji). (**G**) Growth of virulent bacteria *Pma*DG3 in systemic leaves of WT (Col-0) and Dex-inducible complementation lines, three days after infection. The plants were mock treated or immunized as in (**A** and **C**). 30µM Dex plus 0.04% Tween 20 was sprayed on local and systemic leaves 21h before local and distal infection, respectively. (**H**) RT–PCR of the indicated *HyPRP* transcripts in Dex-inducible complementation lines. Leaves were collected 21h after treatment with 30µM Dex (+) or no treatment (-). Expression of *EF1⍺* was used as control. (**I**) Response gain of SAR due to *Pma*DG6 immunization in (**G**). In the graphs showing ‘response gain’, error bars indicate uncertainties in the experiment; in all other graphs error bars show SEM. Different letters above the bars indicate statistically significant difference. Anova, SNK test or Student’s test, p<0.05

We also analyzed the induction of SAR marker PR1 in the uninfected systemic leaves of WT, *azl3, azl13, azl14,* and *elp* mutants after local immunization with *Pma*DG6 or mock treatment. Consistent with the observed bacterial growth (**Figure 7A**), there was increased accumulation of PR1 in systemic leaves of *elp* compared to WT, 72 hours post local *Pma*DG6 infection (**Figure 7E, F**). Moreover, *azl3* and *azl13* mutants showed significantly lower PR1 than WT in systemic leaves at both 48 and 72 hpi (**Figure 7E, F**).

We performed complementation experiments **(Figure 7G)** using the transgenic plants in which GFP fusions were studied in **Figure 3**. The expression of HyPRPs was induced by spraying whole plants with Dex plus 0.04% Tween 20, 21h before infection. In addition to visualizing GFP (**Figure 3**), transgene expression was confirmed by RT-PCR (**Figure 7H**). Dex:AZL3#1 and Dex:AZL13#1 transgenic lines were complemented for SAR and were indistinguishable from WT plants (**Figure 7G, I**). Independent Dex:AZL3#2 and Dex:AZL13#2 lines were also complemented for SAR (**Supplementary Figure S4 B, C**). The Dex:ELP#1 line was completely SAR-deficient, indicating that the dosage of ELP (overexpression) strongly affects SAR (**Figure 7G, I**). The contrasting phenotypes of the mutant (super-SAR) and Dex:ELP#1 line (SAR-deficient) strongly position this particular HyPRP as a negative regulator of SAR.

### HyPRPs are required for ISR

Many HyPRPs are differentially expressed in roots, particularly during the interaction of roots with *P. simiae* WCS417 (**Table 1;** (Stringlis et al., 2018)). Therefore, we tested the role of several HyPRPs with plastid-associated pools in ISR. As shown in **Figure 8**, *azl3, azl13* and *azl14* mutants were unable to mount ISR. In contrast, only WT and *elp* showed significant (and similar) ISR (**Figure 8A, B**). These data indicate that several newly characterized HyPRPs have positive roles in promoting ISR.

**Figure 8.**
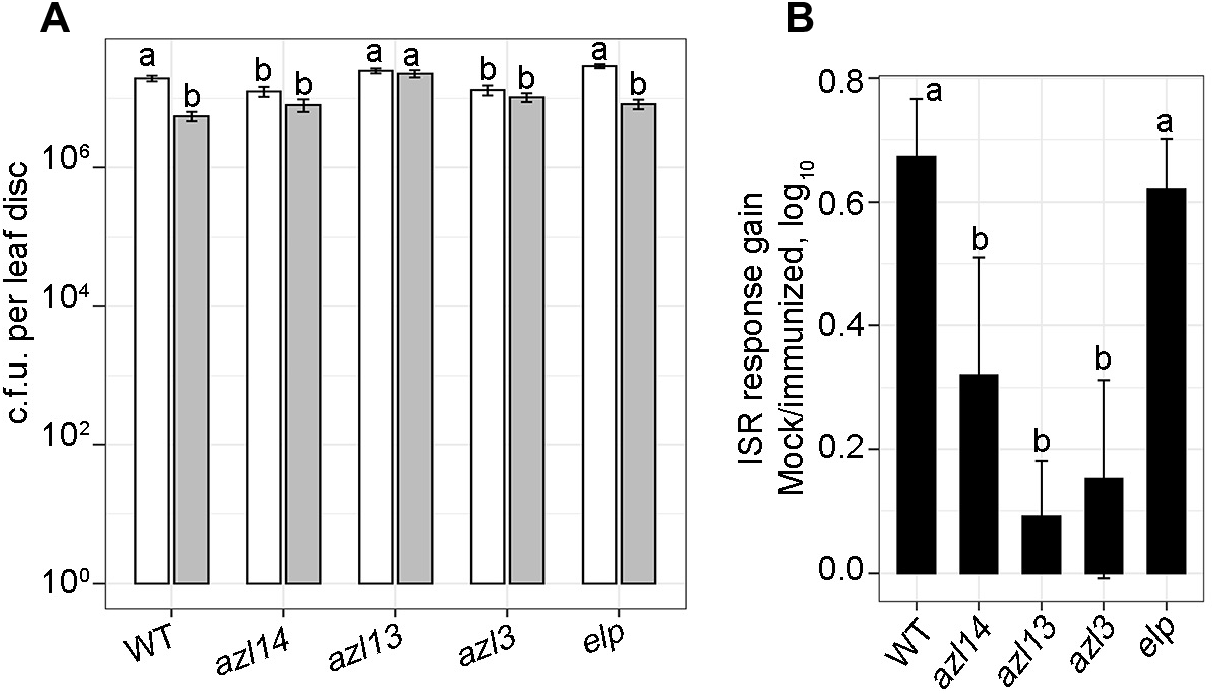
ISR in *HyPRPs* mutants in Arabidopsis. (**A**) ISR was assayed by quantification of virulent bacteria *Pma*DG3 in leaves of plants, three days after infection. The roots of the plants were mock-treated or inoculated with *P. simiae WCS417,* 15-18 days before challenging the leaves with *Pma*DG3. Error bars show SEM. (**B**) Response gain of ISR due to *P. simiae WCS417* immunization in roots of plants in (**A**). Error bars indicate uncertainties in the experiment. The mean c.f.u. of two independent experiments are plotted. Different letters above the bars indicate statistically significant difference by anova, SNK test, p<0.05

### HyPRPs determine colonization, developmental and/or growth-promoting effects of root-associated bacteria

Strain *P. simiae* WCS417, used to stimulate ISR, also promotes plant growth and lateral root formation in Arabidopsis (Haney et al., 2015; Pieterse et al., 2014). Unlike WT, upon treatment of roots with *P. simiae* WCS417, the *azl13* and *azl14* mutants showed no significant increase in biomass or lateral root numbers. The *azl3* mutant also failed to show a biomass increase, but did display increased lateral root numbers (even higher than WT, **Figure 9**). As was seen with the SAR assay, the *elp* mutant was hyper-responsive, showing both biomass and lateral root numbers that exceeded that seen in WT upon *P. simiae* WCS417 inoculation.

**Figure 9.**
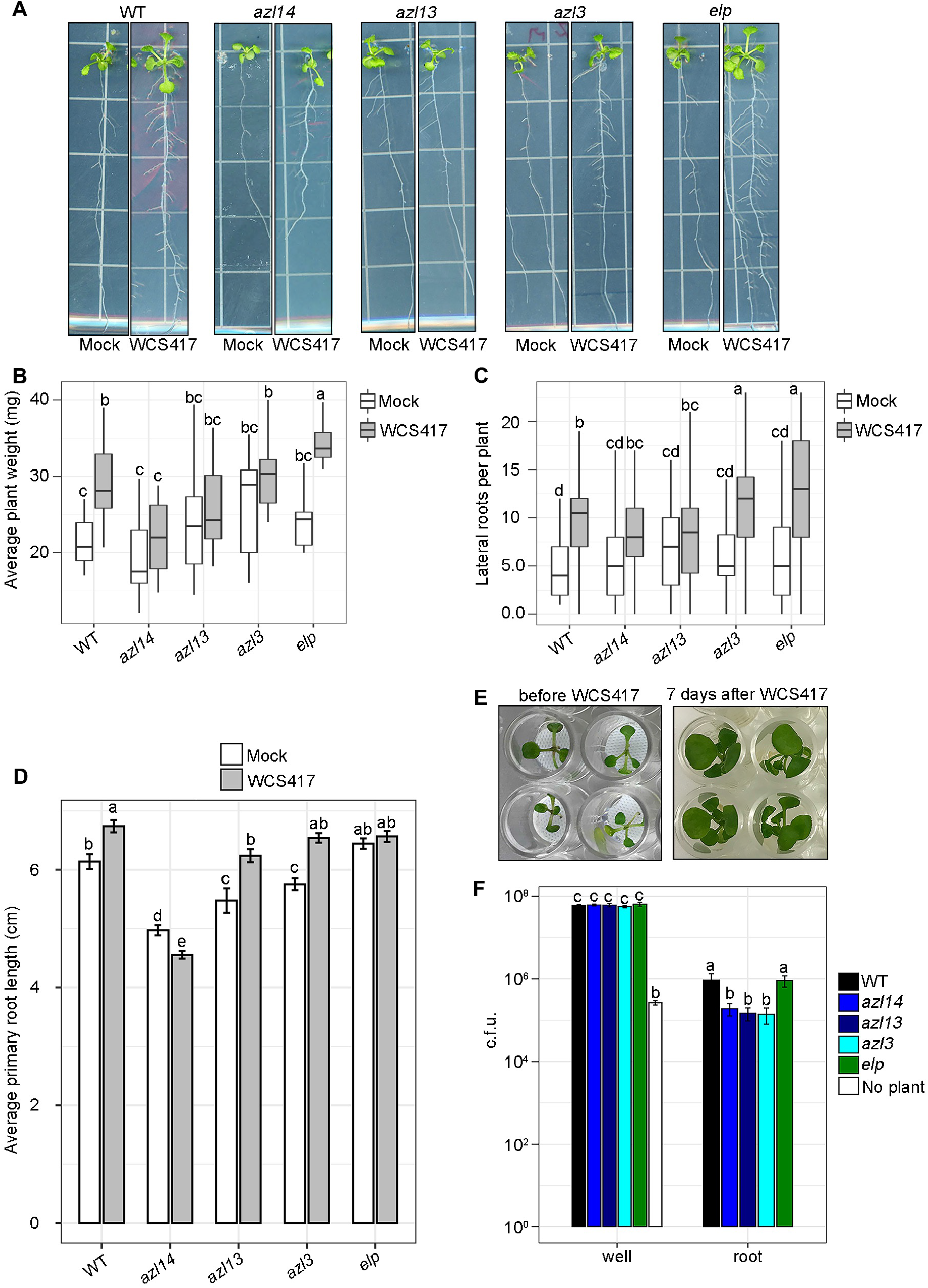
Growth promotion and root-colonization of *HyPRP* mutants by *P. simiae* WCS417. (**A**) Representative images of WT (Col-0) and indicated mutants grown on agar plates without sugar and inoculated with mock or *P. simiae* WCS417 (WCS417). 15 day-old seedlings, 10dpi with WCS417 or mock are shown. (**B**) Fresh weight, (**C**) number of lateral roots and (**D**) primary root length in plants inoculated with mock or WCS417 as in (**A**), n=15-20 plants. Combined data from three independent experiments is plotted. (**E**) Representative images of WT (Col-0) plants grown hydroponically in 48-well plates before (12 day-old) and 7dpi (19 day-old) with WCS417. (**F**) Number of c.f.u. of WCS417 in the wells or attached to the roots of WT and mutants. ‘No plant’ represents the number of c.f.u. of WCS417 in the well-media without any plant present. n=20-24 plants. In all graphs error bars are SEM and different letters above bars indicate statistical differences by anova, SNK test. p<0.05.

We noticed that *azl14* and to a lesser extent *azl3* and *azl13* had shorter primary root lengths than WT in the absence of bacteria (**Figure 9D**). However, *azl3* and *azl13* showed similar increased primary root growth as WT in response to WCS417. In contrast, the *elp* mutant failed to show increased primary root growth and *azl14* showed reduced growth in response to *P. simiae* WCS417.

Differences in *P. simiae-*induced phenotypes may be related to altered levels of root colonization (Haney et al., 2015). Therefore, we analyzed the root bacteria attachment in WT and mutant plants after growing the different genotypes hydroponically in 48-well plates on a Teflon mesh (**Figure 9E**). Seedlings were initially grown with 2% sucrose to avoid differences in root development between genotypes and thus allowing subsequent comparisons of colonization of roots in media without sucrose (see Methods). Interestingly, although there was no significant difference in the bacterial growth (thought to be stimulated by root exudation) in the well media in which plants were growing, *azl3, azl13* and *azl14* mutant roots supported less *P. simiae* colonization compared to WT or *elp* mutant plants (**Figure 9F**).

Altogether, these data suggest that rhizosphere-mediated differences in *P. simiae-*induced responses may partly depend on the host’s ability to support bacterial growth on the roots. Importantly, these results suggest that HyPRPs have varied positive and negative role(s) in the roots including supporting beneficial microbe colonization and impacting root architecture and growth promotion.

## Discussion

Most HyPRP family proteins have SP-like/HD domains at their N termini followed by varying lengths of unstructured PRRs. We previously proposed that the N-terminal bipartite signal formed by SP-like/HD (TMD) + PRR define the subcellular targeting of AZI1 and paralogs to plastids (Cecchini et al., 2015, 2020). By using algorithms (PATS, iTargetP/iChloroP) that recognize the N-terminal signal (SP-like/TMD + PRR/cTP or just PRR/cTP), we predicted with about 80% success the subcellular targeting of many members of the HyPRP family in Arabidopsis. Predictions that were not correct largely correlated with HyPRPs harboring shorter PRRs. Importantly, the PATS algorithm also identified plausible bipartite signals in a large number of Arabidopsis proteins. PATs correctly predicted ≥ 33% of the non-HyPRP proteins that we tested. The non-HyPRP plastid protein STMP6 is among a group of proteins recently proposed to be secreted (Yu et al., 2020). Many proteins in this group, regulated in response to pathogens, are PATS-positive and are thus also good candidates for having a pool that localizes to plastids.

Plastid targeting signals are diverse, consisting of multiple sequence motifs that are distinct or conserved (Bruce, 2001; Lee et al., 2008). Of note is that Pro residues in cTPs mediate efficient translocation of TMD-containing proteins through the chloroplast envelopes (Lee et al., 2018). A Ser/Pro-rich domain targets or facilitate targeting/anchoring of other inner envelope proteins (Küchler et al., 2002; Tripp et al., 2007). Longer PRRs together with the TMDs may similarly help in chloroplast membrane-anchoring of HyPRPs (Cecchini et al., 2020). Charged amino acids (intrinsic charge or those caused by post translational modifications) may play a role in targeting, given their prevalence in PRRs (**Supplementary Table S1**). Considering that AZI1 has a non-cleavable N-terminal signal sequence, these domains in other HyPRPs also are unlikely to be cleaved during their import (Cecchini et al., 2020). This is consistent with the finding that most known proteins in the outer envelope lack a cleavable targeting signal (Hofmann and Theg, 2005). Thus, it is plausible that other plastid-membrane targeted HyPRPs follow the targeting mechanism similar to AZI1, with SP-like/TMD + PRR guiding their plastid association.

We showed that four HyPRPs (AZL2, AZL13, AZL14 and ELP) are OEM plastid proteins. Two HyPRPs (AZL3 and DRN1) were previously suggested to be thylakoid and/or envelope proteins, as evidenced by their presence in proteomics datasets (Peltier et al., 2004; Tomizioli et al., 2014; Kleffmann et al., 2004). While thylakoid samples could be contaminated with plastid envelope proteins, our microscopy of AZL3 in Arabidopsis is consistent with a thylakoid location. This suggests that bipartite signals, depending on their composition, may permit the targeting of proteins to plastid envelopes or the internal plastid thylakoid membrane.

**Figure 10** summarizes the localization patterns of HyPRPs, and by analogy to AZI1/EARLI1, proposes that they are needed for mobilizing signals. The multiple roles of HyPRPs in microbial responses that were found previously (Cecchini et al., 2015, 2019b; Dhar et al., 2020) and herein are shown in **Figure 10B**. ELP is unique among the HyPRPs in how it impacts responses to microbes, as it is a negative regulator of both SAR and growth/development responses to the root-colonizing microbe WCS417. The *elp* mutant shows increased response gain and PR1 accumulation during SAR, but does not affect basal resistance, suggesting increased signaling output to distal tissue. This mutant also has normal root colonization, but has increased plant growth and lateral root numbers relative to WT, further supporting enhanced signaling. However, ISR is not affected in the *elp* mutant, indicating distinct requirements for the developmental responses and ISR. This is consistent with certain ISR-defective mutants showing robust growth and developmental responses when co-cultivated with WCS417 (Zamioudis et al., 2013).

**Figure 10.**
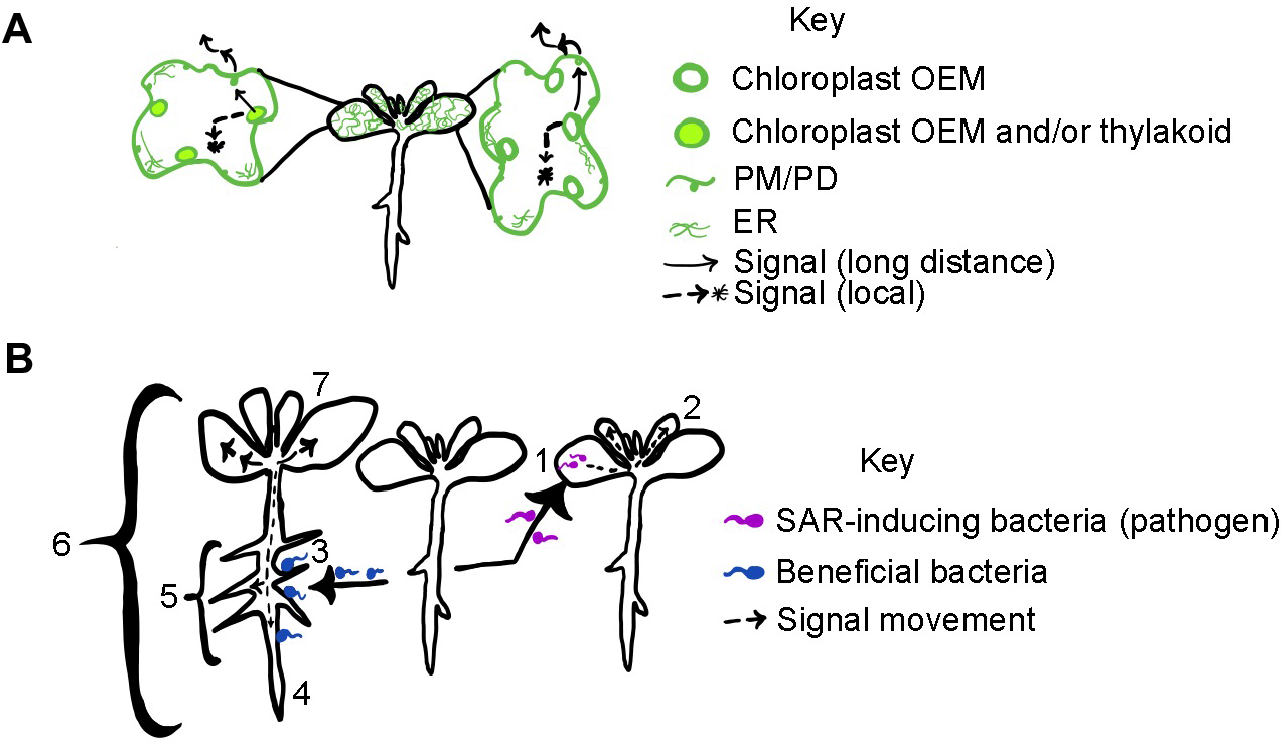
Speculative model and summary of roles of HyPRPs in microbial interactions. (**A**) Cellular localization and proposed roles of HyPRPs in signal mobilization. Green color indicates organelles and structures in cells where HyPRPs are found. Arrows show the potential mobilization of signals by HyPRPs, either within a cell or locally near the infection (dashed line and *) or the long-distance mobilization of signals to distal tissue. Although leaf epidermal cells are depicted, similar localization patterns are expected for root cells. HyPRPs with * have roles in local responses to microbes (root or leaf colonization); most also have roles in longer distance responses as well (see part B). Data from this and previous work (Cecchini et al., 2015, 2020; Peltier et al., 2004; Tomizioli et al., 2014) is summarized. The cell on the left shows HyPRPs that likely locate to plastid thylakoids and/or OEMs as well as ER and PM/PD. HyPRPs that fit the left pattern, AZL3* and DRN1*, have positive roles. The cell on the right shows HyPRPs that locate to plastid OEMs as well as ER and/or PM/PDs. HyPRPs that fit the right pattern have positive roles (AZI1, EARLI1, AZL2, AZL13, and AZL14), a mainly negative role (ELP) or have unknown roles (CWLP, DHyPRP, AZI3, AZI4) in microbial responses. (**B**) Schematic of the steps at the whole plant level that are affected by one or more HyPRPs: 1, leaf colonization by pathogen (DRN1, positive role); 2, increased immunity to distal infections after an immunizing infection at site 1 (SAR: AZI1, EARLI1,AZL2, AZL3, AZL13, positive roles; ELP, negative role); 3, growth of beneficial bacteria in association with roots (AZL3, AZL13, AZL14, positive roles); 4, stimulation of primary root growth in response to beneficial bacteria colonization of the root (ELP, AZL14, positive roles); 5, stimulation of lateral roots in response to beneficial bacteria colonization of the root (AZL13, AZL14, positive roles; ELP, negative role); 6, stimulation of whole plant growth by root colonization by beneficial bacteria (AZL3, AZL13, AZL14, positive roles; ELP, negative role); 7, increased immunity of aerial tissue (ISR after root colonization by beneficial bacteria: AZI1, EARLI1, AZL3, AZL13, AZL14, positive roles).

Some mutants in this study (*azl3* and *azl13*) are affected in both SAR and ISR, similar to the *azi1* and *earli1* mutants of HyPRPs (Jung et al., 2009; Cecchini et al., 2015). For the SAR phenotypes, the defects are linked to long distance signaling, since the growth of the immunizing pathogen is normal. However, the *azl3*, *azl13* and *azl14* mutants show possible local signaling defects evidenced by lower colonization of roots with WCS417 relative to that seen in WT. It is possible that other root-related defects in these mutants arise from low root colonization. A certain threshold of rhizobacterial colonization may be needed to induce the downstream root development, plant growth and ISR phenotypes. However, *azl3* still shows lateral root induction, indicating that there is enough colonization to cause signaling for this phenotypic output. Lateral root induction in *azl3* may occur due to the presence of other HyPRPs (AZL13, AZL14, AZI1 and/or EARLI1 (Cecchini et al., 2019b)). This would implicate other HyPRPs in regulating both colonization as well as the downstream signaling for lateral root induction.

It seems possible that different phenotypic outputs (growth promotion, ISR and lateral root number increases) require different thresholds of bacterial colonization of the roots. Alternatively, all outputs may require the same threshold, but the certain HyPRPs may be differentially needed for multiple steps: colonization and downstream signaling to cause growth, ISR and/or lateral root production. A genome-wide association mapping study (Wintermans et al., 2016) identified different markers that were independently associated with shoot growth promotion, lateral root formation and primary root length changes in response to WCS417. This finding supports the idea that there are separate pathways (or branches of a pathway) that independently regulate the different phenotypic outputs. Interestingly, two HyPRPs (DEG27, AZL2) are within 5-10 KB of a marker associated with regulating the lateral root changes.

Despite having positive roles in regulating interactions with/responses to microbes, transcript levels of many HyPRPs are down regulated by microbe-related signals. This could reflect negative feedback to modulate signaling. However, in plants with constitutive defenses (ACD6-1HA) the transcript level of AZL3 is low, while the plastid membrane-associated protein level is high relative to WT. This suggests that there is complexity to the regulation of HyPRPs. Some HyPRPs may become stabilized by post translational modification, since they have kinase motifs (SP/TP, **Supplementary Table S1**). AZI1’s PRR is a kinase substrate *in vitro* (Pitzschke et al., 2014).

HyPRPs are similar and they locate to several types of membranes. This could permit the formation of membrane contact sites and complexes with other HyPRPs/LTPs in different places. Multiple contact sites between plastids, ER and PM, comprising of LTPs, are the sites of exchange of small molecule signals (Andersson et al., 2007; Li et al., 2020; Toulmay and Prinz, 2011). HyPRPs are likely enriched in these sites for the mobilization/transport of immune and/or developmental and growth signals. Supporting this, AZI1 and EARLI1 can interact and mobilize the SAR signal AZA, which also can trigger AZI1-/EARLI1-dependent root architecture changes (Cecchini et al., 2019b, 2015). Moreover, because AZI1/EARLI1 physically interact with non-HyPRP LTP superfamily member (i.e. DIR1) in membrane contact sites to modulate systemic responses (Cecchini et al., 2015; Yu et al., 2013), it is also possible that other positive regulatory HyPRPs similarly form active homo or hetero-oligomeric LTP complexes at the key sites/structures for transmission of developmental, growth and/or defense signals to regulate systemic defense responses. The presence of so many HyPRPs may allow the movement of signals at different rates (to reach specific thresholds) or enable movement of various signals that require different membrane-paths to move. For example, to reach the xylem to move signals aboveground may require the use of different contact sites compared to signals reaching parenchyma for lateral root stimulation. Thus, different signals (or the amount of signal moved) could affect and explain different specific phenotypes.

## Methods

### Plants, bacteria and plasmids

All the *Arabidopsis thaliana* plants used were Columbia-0 (Col-0) ecotype. The plants used for infection were grown in soil (Berger, BM1:BM2; 50:50 mix) or Jiffy-7^®^ peat pellets (for ISR, Hummert International, # 14-23700) at 12h day/night cycle as previously described (Cecchini et al., 2015). The following Arabidopsis accessions were obtained from ABRC: *At1g12090* (*elp*, SALK_147582C; *elp-2,* SAIL_532_C09), *At2g10940* (*azl3*, SALK_083118C), *At5g46890* (*azl13*, SALK_065132), *At5g46900* (*azl14*, SALK_096750C). *At1g62510 AZL2-AS* line was described previously (Jülke and Ludwig-Müller, 2015). The transgenic line ACD6-1HA (Lu et al., 2005) used for proteomic analysis was grown at 16h day/8h night cycle. pt-gk (cTP-GFP) transgenic line from ABRC (CS16266) was used as a chloroplast marker. Arabidopsis HyPRP complementation lines were generated by floral dip method using *Agrobacterium tumefaciens* C58C1 suspensions (Clough and Bent, 1998). *N. benthamiana* used for localization studies were grown in soil at 24°C under 16h light/8h night regime. For *A. tumefaciens*-mediated transient assays, 4–5-week-old *N. benthamiana* plants were used.

Bacterial strains used for infections or colonization were virulent *Pseudomonas cannabina pv. alisalensis* (formerly *Pseudomonas syringae pv. maculicola* strain ES4326) carrying an empty vector (*Pma*DG3), the avirulent isogenic strain carrying avrRpt2 (*Pma*DG6) and *Pseudomonas simiae* WCS417 (formerly *Pseudomonas fluorescens* WCS417r).

The full-length coding sequences or the truncated versions of HyPRPs were amplified by PCR and cloned by Gateway procedure into Dexamethasone (Dex)-inducible plant expression vectors pBAV150 (C-terminal GFP tag) or pBAV154 (C-terminal HA tag) (Cecchini et al., 2015). Full-length coding sequences of HyPRPs in pBAV150 vector were used to transform Arabidopsis to generate complementation lines. To generate ΔPRR variants, proper fragments were generated with overlapping primers and linked by PCR. ΔLTP variants were generated using non-overlapping primers in PCR. All the primers and constructs used in this study are listed in (**Supplementary Table S2**).

### *In silico* analysis of subcellular targeting

For bipartite plastid targeting signal prediction, we analyzed the full-length sequences of Arabidopsis proteome (ATpepTAIR10) by using PATS software (http://modlabcadd.ethz.ch/software/pats/) or an iterative method (iTargetP/iChloroP). In the iterative method, N-terminal SP-like regions as determined by TargetP (http://www.cbs.dtu.dk/services/TargetP-1.1/) or SignalP (http://www.cbs.dtu.dk/services/SignalP/), varying between 22 and 27 amino acids (except for AZI7 with 30 amino acids as SP-like) were removed *in silico* and the remaining sequences were analyzed by TargetP/ChloroP to determine the presence of chloroplast transit peptide following the SP-like region.

The GO analysis was done as described (Bonnot et al., 2019). The list of representative enriched GO terms was obtained using Panther (https://www.arabidopsis.org/tools/go_term_enrichment.jsp) and REVIGO (http://revigo.irb.hr) tools.

### Confocal Imaging

9-12 day-old Arabidopsis seedlings or 4-week old *N. benthamiana* leaves expressing the different HyPRP versions fused at C-terminal with GFP were visualized by confocal microscopy. Zeiss LSM710 and LSM800 laser scanning confocal microscopes were used to visualize GFP fluorescence (excitation = 488 nm, emission = 509 nm) and chlorophyll autofluorescence (excitation = 631 nm, emission = 647 nm). LD C-Apochromat 40x/1.1 W Korr (LSM710) and EC Plan-Neofluor 40x/1.3 (LSM800) objectives were used for taking images. The images are optical sections captured at 1024 x 1024 pixels scanning resolution in maximum speed mode. Fluorescence of GFP and chlorophyll was acquired in sequential acquisition mode. Plant tissues were mounted in perfluorodecalin (Strem Chemicals, Inc.) for optical enhancement. Images were processed by ImageJ (Fiji) and Adobe photoshop software.

### Fractionation

*N. benthamiana* leaves, transiently transformed with *A. tumefaciens* harboring different constructs, were used to isolate chloroplasts as described in (Cecchini et al., 2015) with minor modifications. Briefly, leaves (0.5-1g) were homogenized in XpI buffer (0.33 M sorbitol, 50 mM HEPES pH 7.5, 2 mM EDTA, 1 mM MgCl_2_, 0.25% BSA, 0.1% sodium ascorbate and protease inhibitors) using polytron (Kinematica) and filtered through two layers of Miracloth (Calbiochem, # 475855). The filtrate was centrifuged at 5000 rpm for 5 minutes and the pellet was resuspended in 0.4ml of XpI buffer. The resuspended pellet was loaded on Percoll (GE, # 17089101) gradient (80% and 40%) and centrifuged at 13,000 rpm to isolate the intact chloroplasts. The recovered intact chloroplasts at the interphase were resuspended in 0.5ml XpI buffer and centrifuged at 3000 rpm for 5 minutes to remove the Percoll impurities. The purity of the chloroplast fractions was checked using specific markers by immunoblotting.

To obtain enriched membrane and soluble fractions, the intact chloroplasts were partitioned into membrane (pellet) and soluble fractions by centrifugation. To do this, the plastids were resuspended in protease free ice-cold water containing protease inhibitor cocktail (Thermo Scientific, #A32965). For complete lysis, the chloroplasts were incubated on ice for 45-60 min followed by incubation at −80°C O/N. The samples were then thawed on ice and centrifuged at 13,000 rpm. The soluble fraction was recovered, and the pellet was resuspended in 1XPBS (+1% SDS and protease inhibitor cocktail).

### Thermolysin assay

Proteolysis by thermolysin was performed as described in (Cecchini et al., 2015). Briefly, intact chloroplast fractions were resuspended in 1X import buffer (0.33 M sorbitol, 50 mM HEPES, pH 8.0) and treated with thermolysin (to a final concentration of 0.1mg/ml). The reaction was quenched by adding 10mM EDTA. The chloroplasts were carefully separated on 40% Percoll cushion.

### Immunoblotting

Equal amounts of total proteins were separated on 12% SDS-PAGE and transferred to PVDF membranes (Millipore, #IPVH00010). The concentration of proteins was determined by Bradford assay. The following primary antibodies were used in this study: PR1 antibody (Agrisera, #AS10687; 1:2,500), GFP monoclonal antibody (Takara, # 632375; 1:2,000), Bip2 antibody (Agrisera, #AS09481; 1:4,000), Tic110 antibody (a gift from Masato Nakai; 1:3,000), chloroplast Hsp70 (a gift from Thomas Leustek; 1:12,000), Toc75 (Agrisera, #AS08351, 1:3000). Horseradish peroxidase-conjugated anti-mouse (Invitrogen, #SA1-100; 1:4,000), anti-guinea pig (Sigma, #A5545; 1:100,000), or anti-rabbit (Thermo Scientific, #32460: 1:5,00) secondary antibodies were used. The bands were detected by using SuperSignal pico/femto chemiluminescence kits (Thermo Scientific; #’s1859022/3 and 1859674/5).

### Transcript analysis

Total RNA was isolated with the RNeasy Plant Mini Kit (Qiagen) and cDNA was synthesized using was the Reverse Transcriptase SuperScript III and Oligo (dT)_20_ primers (ThermoScientific) according to the manufacturer’s instructions. *HyPRP* expression (in WT/ACD6-1HA) was analyzed by quantitative RT-PCR using SYBR Premix Ex Taq (Takara) in a Bio-Rad Real-Time CFX96TM C1000 system (Bio-Rad) according to the manufacturer’s instructions. For Dex:AZL lines, relative transcript levels were checked by RT-PCR (optimized for cycle #s). EF1α was used as a reference for normalization.

### Pathogen growth bioassays

The evaluation of bacterial pathogen growth was caried as described (Jung et al., 2009). Briefly, 26-28 day-old wild-type and *HyPRP* mutant plants were syringe-infiltrated with *Pma*DG3 or *Pma*DG6 (OD_600_=0.0001). Growth was quantified using eight equal sized leaf discs from different plants 3 days after bacterial inoculation. All graphs of colony forming units (cfus) were plotted as log_10_ values.

For SAR assays (Jiang et al., 2021), local leaves of 26–28-day old plants were syringe-infiltrated with mock (10mM MgSO_4_) or *Pma*DG6/avrRpt2 (OD_600_ = 0.01). Two days after the primary infection, systemic leaves were infiltrated with *Pma*DG3 (OD_600_= 0.0001). The local leaves were removed from the plants prior to the secondary challenge and the plants were covered with plastic domes following the secondary challenge. Three days post-secondary challenge, equal sized leaf discs were collected and ground in 10mM MgSO_4_. The ISR assay was performed exactly as described (Cecchini et al., 2019a) with *Pma*DG3 OD_600_ of 0.0001-0.0002. The response gain of SAR or ISR was calculated as described (Jiang et al., 2021).

### Plant growth promotion assays of whole plants and roots

Arabidopsis were grown vertically at ∼20°C with 16h day/8h night on small square plates containing MS media (GibcoBRL, Life Technologies) without sugar as previously described (Haney et al., 2015). Briefly, ten seeds were germinated on plates and five days after plating, seedlings were thinned to five seedlings per plate and roots inoculated with 1µl 10mM MgSO_4_ (Mock) or *Pseudomonas simiae* WCS417 (OD_600_=0.001). After 10 days, scanned (Epson) plant images were used to count lateral roots and measure primary root lengths. Subsequently the fresh weights were determined.

### Root colonization assay

Root colonization assay was performed in 48-well plates as described (Haney et al., 2015). 250μl MS media (Sigma, #M5519-10L) containing 2% sucrose was added to the wells of 48-well tissue cultures plates (BD Falcon). Round Teflon mesh disks (McMaster-Carr #1100t41) were sterilized by autoclaving and placed into the wells. A single surface sterilized and imbibed seed was placed at the center of each disc. Seedlings were subsequently grown at ∼20°C with 16 h day/8 h dark for 10 days. The MS media was replaced with 270 µl of fresh MS media without sugar. Two days after the media change, wells were inoculated with 30 μl (final OD_600_ = 0.00002) of *P. simiae* WCS417 suspended in water. After 7 days, seedlings were removed, and roots and well media analyzed for bacterial amount (six seedlings were combined per data point). The root extracts prepared by grinding and the corresponding well media were plated by serial dilution to determine cfus.

### LC-MS/MS

Approximately 15g Arabidopsis shoots of WT and ACD6-1HA were homogenized and the chloroplasts separated on Percoll gradients (80% and 40%). The total plastid membrane fraction obtained after centrifugation was used for LC-MS/MS. Total plastid membrane proteins were analyzed by Q-Exactive Orbitrap mass spectrometer. Relative quantification was performed using Maxquant software. The raw data was searched against the *Arabidopsis thaliana* database, as well as reversed protein sequences, and common contaminants. The search results were filtered to be within 1% false discovery rate and label-free quantification using normalized protein abundances was performed to assess differential protein expression between WT and the transgenic expressing *ACD6-1HA*.

### Plotting and statistical analysis

The graphs were plotted in R studio (v.3.6.1/4.1) using ggplot2 package (v3.3.2) and all statistics were done using agricolae package (v1.3.3).

## Acknowledgments

This work was supported by NSF grant IOS-1456904 to J.T.G. Z.Z.B. was supported by a fellowship from SERB/IUSSTF. At the University of Chicago, we thank Dr. Atif Khan for help with R programming, Dr. Joanna Jelenska for experimental advice, Professor Stephen Kron and Donald Wolfgeher for assistance with the proteomics experiment, Greenberg lab members for reading and commenting on the manuscript and Dequantarius J. Speed for useful discussions. We thank Professor Jutta Ludwig-Mueller, Technische Universität Dresden for providing the AZL2-AS lines and Dr. Jennifer Morrell-Falvey at Oak Ridge National Laboratory for helpful comments on the manuscript.

## Author contributions

J.T.G., N.M.C. and Z.Z.B. designed the experiments. Z.Z.B, N.M.C, A.T.S., C.T.H., R.C.F., and E.A. performed experiments. Z.Z.B, N.M.C., A.T.S. and J.T.G. performed bioinformatic analysis or data mining. Z.Z.B, N.M.C and J.T.G. wrote the manuscript.

## Supplemental data

**Supplementary Figure S1.** Over-represented GO terms for biological processes in PATS plastid predicted proteins, location of *HyPRP*s in Arabidopsis genome and GFP cleavage in DRN1.

**Supplementary Figure S2.** ClustalW alignment of Arabidopsis HyPRPs.

**Supplementary Figure S3**. Local disease resistance of *HyPRP* mutants against *Pseudomonas syringae* pv. *maculicola strains*.

**Supplementary Figure S4.** SAR in *elp-2* allelic mutant and Dex-inducible complementation lines.

**Supplementary Dataset S1**. List of all PATS positive proteins of Arabidopsis.

**Supplementary Table S1**. CPRs, PRRs, WWH scores, experimental and predicted plastid targeting of Arabidopsis HyPRPs.

**Supplementary Table S2**. List of primers and constructs used in this study.

## Supplementary Figure Legends

**Supplementary Figure S1. Over-represented GO terms for biological processes in PATS plastid predicted proteins, location of *HyPRP*s in Arabidopsis genome and GFP cleavage in DRN1.** (**A**) The most representative and significant GO terms sorted by fold enrichment are plotted. The size of dot indicates the number of genes associated with the process and the color of dot indicates the significance of the enrichment {- log_10_(FDR-corrected *P*-values)}. The vertical dashed line represents a fold enrichment of 1. (**B**) Location of HyPRP genes depicted in Arabidopsis genome, made with the TAIR chromosome map tool (https://www.arabidopsis.org/jsp/ChromosomeMap/tool.jsp). (**C**) Confocal micrograph of epidermal cells and immunoblotting of total (T) fraction of *N. benthamiana* leaves transiently expressing DRN1-GFP showing cleavage of GFP. White arrowhead shows cytoplasm and white arrow indicates nucleus. Scale bar is 20µm. Asterisk shows the free GFP detected by anti-GFP antibody. rbcL is the non-specific rubisco large subunit.

**Supplementary Figure S2. ClustalW alignment of Arabidopsis HyPRPs.** The bars above the alignment indicate the putative SP-like or HD, PRR and 8CM/LTP domain. Black and grey boxes indicate identical or similar sequences, respectively. Red asterisks show the eight conserved Cys residues. AZL4 and AZL8 were excluded due to relatively larger sizes. ClustalW and BoxShade (Expasy tools) were used for the alignment.

**Supplementary Figure S3**. **Local disease resistance of *HyPRP* mutants against *Pseudomonas syringae* pv. *maculicola strains*.** Wild-type (Col-0) and indicated *HyPRP* mutants were infected with (**A**) *Pma*DG3 (virulent strain) and (**B**) *PmaDG6* containing *avrRpt2* (SAR-inducing strain) at OD_600_=0.0001 and growth measured 3 days post infiltration. Error bars indicate SEM. Figures represent the combined data from two independent experiments with n=7-8 for each replicate. Anova (SNK test) showed no statistically significant difference between WT and *HyPRP* mutants.

**Supplementary Figure S4. SAR in *elp-2* allelic mutant and Dex-inducible complementation lines.** SAR assay showing the growth of virulent bacteria *Pma*DG3 in systemic leaves of (**A**) *elp-2* mutants and (**B**) Dex-inducible complementation lines (#2), three days after infection. The plants were immunized in local leaves with avirulent *Pma*DG6 or mock treated 2 days prior to secondary challenge. The mean c.f.u. of one experiment is shown. (**C**) RT–PCR of the indicated Dex line (#2). Leaves were collected 21h after treatment with 30µM Dex plus 0.04% Tween 20 (+) or no treatment (-). Expression of *EF1⍺* was used as control. In (**A**, **B**), error bars show SEM. Different letters above the bars indicate statistically significant difference by anova, SNK test, p<0.05.

